# Convergent evolution of multiple mutations improves the viral fitness of SARS-CoV-2 variants by balancing positive and negative selection

**DOI:** 10.1101/2021.12.23.474050

**Authors:** Vaibhav Upadhyay, Casey Patrick, Alexandra Lucas, Krishna M.G. Mallela

## Abstract

Multiple mutations have been seen to undergo convergent evolution in SARS-CoV-2 variants of concern. One such evolution occurs in Beta, Gamma, and Omicron variants at three amino acid positions K417, E484, and N501 in the receptor binding domain of the spike protein. We examined the physical mechanisms underlying the convergent evolution of three mutations K417T/E484K/N501Y by delineating the individual and collective effects of mutations on binding to angiotensin converting enzyme 2 receptor, immune escape from neutralizing antibodies, protein stability and expression. Our results show that each mutation serves a distinct function that improves virus fitness supporting its positive selection, even though individual mutations have deleterious effects that make them prone to negative selection. Compared to the wild-type, K417T escapes Class 1 antibodies, has increased stability and expression; however, it has decreased receptor binding. E484K escapes Class 2 antibodies; however, it has decreased receptor binding, stability and expression. N501Y increases receptor binding; however, has decreased stability and expression. When these mutations come together, the deleterious effects are mitigated due to the presence of compensatory effects. Triple mutant K417T/E484K/N501Y has increased receptor binding, escapes both Class 1 and Class 2 antibodies, and has similar stability and expression as that of the wild-type. These results show the implications of presence of multiple mutations on virus evolution that enhance viral fitness on different fronts by balancing both positive and negative selection and improves the chances of selection of mutations together.

## INTRODUCTION

SARS-CoV-2 pandemic has disrupted the lives of millions of people around the globe for the past 2 years with enormous damage to human life and world economy. Till February 2022, the virus has infected more than 376 million people and claimed about 6 million lives (https://covid19.who.int/). The emergence of the virus variants has contributed to prolonged pandemic even after effective vaccines, therapeutic antibodies and antivirals have been developed within a year of the pandemic.^*1–10*^ Emergence of the variants has caused waves of infection throughout the world, with each wave majorly contributed by a particular variant. Some of the variants are more dangerous than others and have been termed as the variants of concern (VOC). Till date, five variants have been designated as VOCs by WHO (https://www.who.int/en/activities/tracking-SARS-CoV-2-variants/). Apart from these, numerous other variants have emerged in different parts of the world, some of which have been termed as variants of interest. These variants have raised concerns as the recent data has shown decreased efficacy of vaccines and therapeutic antibodies particularly against VOCs.^*11–21*^ New vaccines and therapeutics will be needed to counter the threats posed by variants. It is thus necessary to understand the properties of the viral variants to be able to come up with more informed approaches for vaccine and therapeutics design.

Among the mutations in the viral genome, the mutations in the receptor binding domain (RBD) are of particular interest as RBD is immunodominant and target of major neutralizing antibody response against the virus.^*22*^ Also, RBD is functionally important as it allows virus to interact with ACE2 receptor on human epithelial cells.^*23–25*^ Thus, RBD mutations are likely to impact the antibody escape potential and ACE2 interaction, thereby influencing virus infectivity and transmission. All of the VOCs, other than Alpha, contain multiple RBD mutations. Presence of multiple mutations can allow virus to acquire advantageous traits which would otherwise not be selected in the case of a single mutation. For instance, a beneficial mutation in terms of antibody escape can have deleterious effect on protein stability or ACE2 binding thereby compromising virus fitness. Such a mutation would not be strongly selected, but in the presence of a compensatory mutation that can improve stability or ACE2 binding, the otherwise deleterious mutation that confers antibody resistance would be strongly selected. Thus, the variants that contain multiple mutations pose a greater threat as they are likely to have greater fitness and better adapted to survive in the host. This makes it important to study the evolution of viral variants that contain multiple mutations and delineate the role of each mutation in the selection of that variant.

Many variants have multiple mutations that are co-evolving. One particular triple mutation that can be consistently found in multiples variants include mutations at K417, E484, and N501 positions in the amino acid sequence of the spike protein RBD. The three mutations are in the protein structural regions that interact with ACE2 and also with major neutralizing antibodies, and hence are likely to impact ACE2 binding as well as antibody binding interactions. In this study, we examined the individual and collective effect of the three RBD mutations K417T/E484K/N501Y on expression, protein stability, ACE2 binding and immune escape potential against the common neutralizing antibodies. To achieve this, we created single, double and triple mutants of RBD. This approach allowed us to determine which mutations offer improved viral fitness with respect to a particular physical parameter, and if there is an association between mutations, synergistic or antagonistic, that determines the selection of mutations together.

## RESULTS

### K417T mutation increases while E484K mutation decreases protein expression

To understand the effects of mutations on expression, stability, ACE2 binding and antibody escape, we created different RBD mutants comprising of single mutations (K417T, E484K, N501Y), double mutations (K417T/E484K, K417T/N501Y, E484K /N501Y) and triple mutations (K417T/E484K/N501Y). The unmutated, wild-type (WT) RBD and the seven mutants were expressed in Expi293 cells which are modified human embryonic kidney (HEK) cells. These proteins were expressed as secretory proteins, which means they traverse through the secretory pathway in human cells and are subject to the endoplasmic reticulum (ER) quality control, which allows well folded proteins to secrete out of the cells while the misfolded and aggregation prone proteins are recognized and degraded through the ER degradation pathway. The protein expression levels can impact the overall virus yield as higher protein expression means greater viral packaging efficiency.^*26*^ The mutations were found to affect the levels of protein that was secreted out of the cells (Figure 1). K417T mutation increased the levels of secretory protein by 70% relative to the WT protein, while E484K mutation decreased the expression level by 60%. N501Y mutation also decreased the expression by 20%. The effect of these mutations was additive in the case of double mutants (with K417T mutation) and the triple mutant studied.

**Figure 1.**
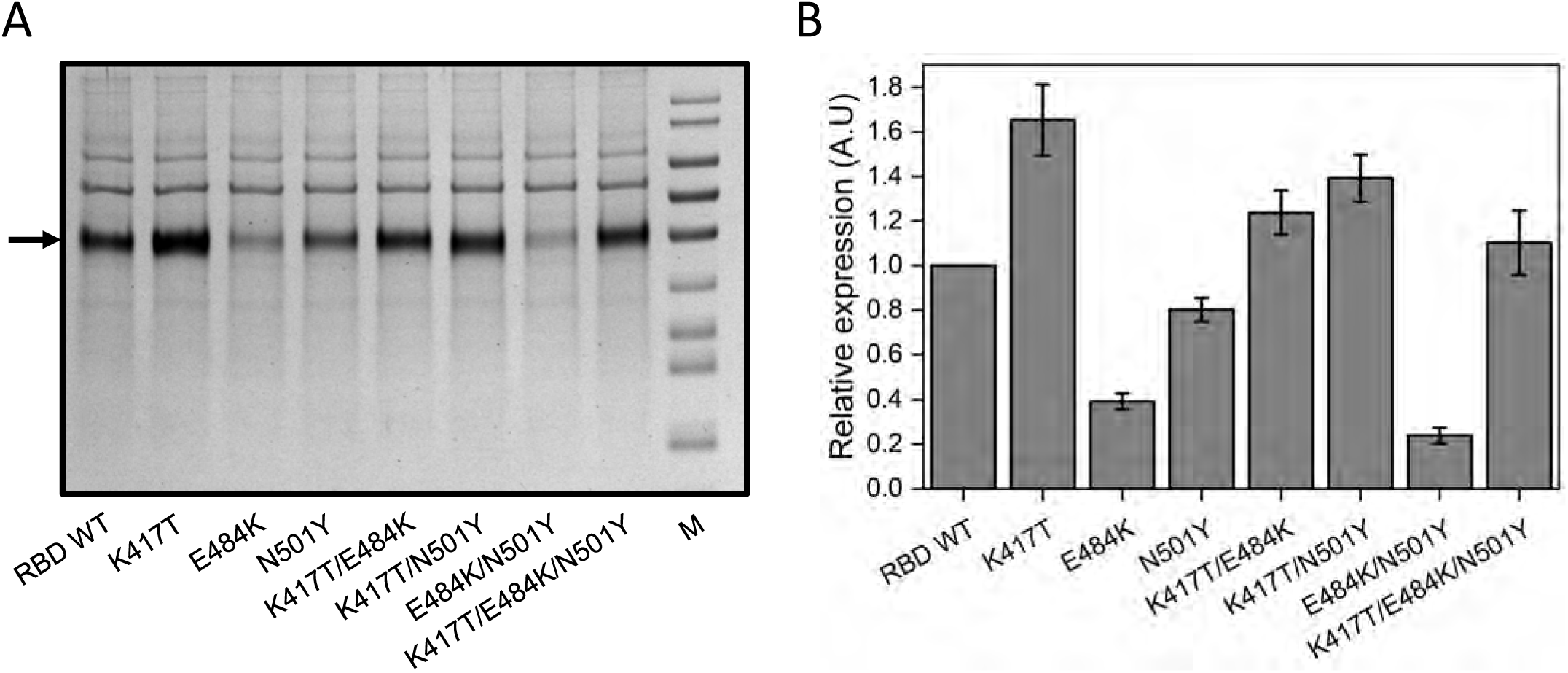
Comparative analysis of protein expression of wild-type RBD and its mutants using SDS-PAGE. (A) SDS-PAGE showing relative amounts of expressed RBD and mutants. M represents molecular weight markers (from top to bottom: 180, 130, 100, 70, 55, 40, 35, 25 and 15 kDa). The arrow indicates the position of the band of interest. (B) Quantification of band intensities shown with arrow in (A) relative to expression of WT RBD.

Mutants that carried K417T mutation had higher expression. K417T/N501Y increased expression by 40%, whereas K417T/E484K and K417T/E484K/N501Y increased expression by 10%. The presence of two deleterious mutations E484K and N501Y together had a negative cooperative effect on the expression of the double mutant E484K/N501Y, which had 80% less expression and could not be purified in high yield. Further studies were therefore done on the WT protein and six mutant proteins excluding the double mutant E484K/N501Y. All proteins used for further studies were purified to homogeneity (Figure 2A). These protein expression results suggest that K417T may provide fitness advantage to the virus by providing more protein for virus assembly and thereby increasing virus transmission.

**Figure 2.**
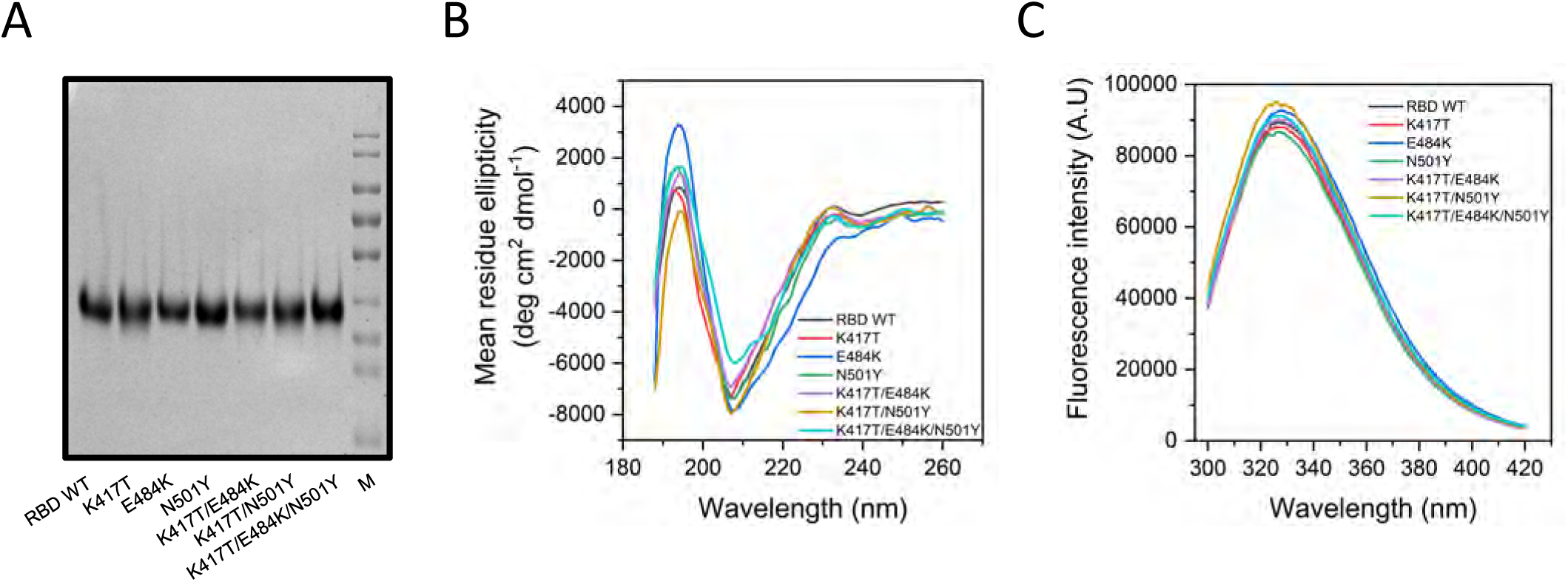
Structural characterization of WT RBD and its mutants. (A) Purified RBD and other mutants. M represents molecular weight markers (from top to bottom: 180, 130, 100, 70, 55, 40, 35, 25 and 15 kDa). (C) Comparison of secondary structure of WT RBD and its mutants using far-UV CD spectra. (D) Comparison of tertiary structure of WT RBD and its mutants using fluorescence emission spectra.

### None of the three RBD mutations affect the global protein structure

Most single amino acid mutations in proteins that have been characterized are in general neutral and do not bring a significant change to protein structure. Other mutations may be deleterious or bring an advantage to the organismal fitness. The deleterious mutations are generally not selected unless a selection pressure like immune response acts on them. Advantageous mutations on the other hand are selected. We investigated the effect of mutations on the secondary and tertiary structure of RBD using circular dichroism (CD) and fluorescence spectroscopy to determine if any of the mutations affect the global protein structure (Figures 2B and 2C). The results suggest that the global secondary as well as tertiary structure is not affected by the mutations, even though the three mutations are non-conservative where K417T mutation changes a charged amino acid to an uncharged amino acid, E484K changes a negatively charged amino acid to a positively charged amino acid, and N501Y changes a polar amino acid to an aromatic amino acid.

### K417T mutation stabilizes RBD while E484K mutation destabilizes RBD

Protein stability is an important parameter that impacts the evolution of viral proteins.^*27*^ More stable proteins can withstand higher number of deleterious mutations and thus tend to persist for longer time in the viral pool.^*28*^ We measured the stability of the WT and mutated RBDs using thermal and chemical denaturation melts. The thermal denaturation melts were recorded using far-UV CD spectroscopy. Figures 3A-3G show the representative melts. Thermal melts were not completely reversible and are thus not indicative of the equilibrium thermodynamic stability of the proteins. Therefore, thermal melts can only be used to compare the mid-point of thermal denaturation (T_m_) and are good measure of apparent stability of proteins.^*29*^ Table 1 lists the T_m_ values along with errors determined independently from three different batches of protein expression. The T_m_ of WT RBD was 56.1 ± 0.7 °C. N501Y mutation did not have a significant impact on the T_m_ value compared to the WT protein, with slightly decreased T_m_ value of 55.1 ± 0.3 °C. K417T mutation increased the thermal stability of the proteins. Mutants carrying K417T mutation have higher T_m_ values compared to the WT protein with T_m_ values of 59.3 ± 0.2, 57.2 ± 0.5, and 59.1 ± 0.2 for K417T, K417T/E484K, and K417T/N501Y, respectively. E484K mutation on the other hand decreased the thermal stability of the protein, with the single site mutant having a T_m_ value of 52.3 ± 0.5 °C. The effect of mutations on thermal stability was also found to be additive because of which the mutants that carry double or triple mutations have T_m_ values that are approximately the average of the T_m_ values of individual mutants. The triple mutant K417T/E484K/N501Y has similar thermal stability as that of the WT (Table 1).

**Figure 3.**
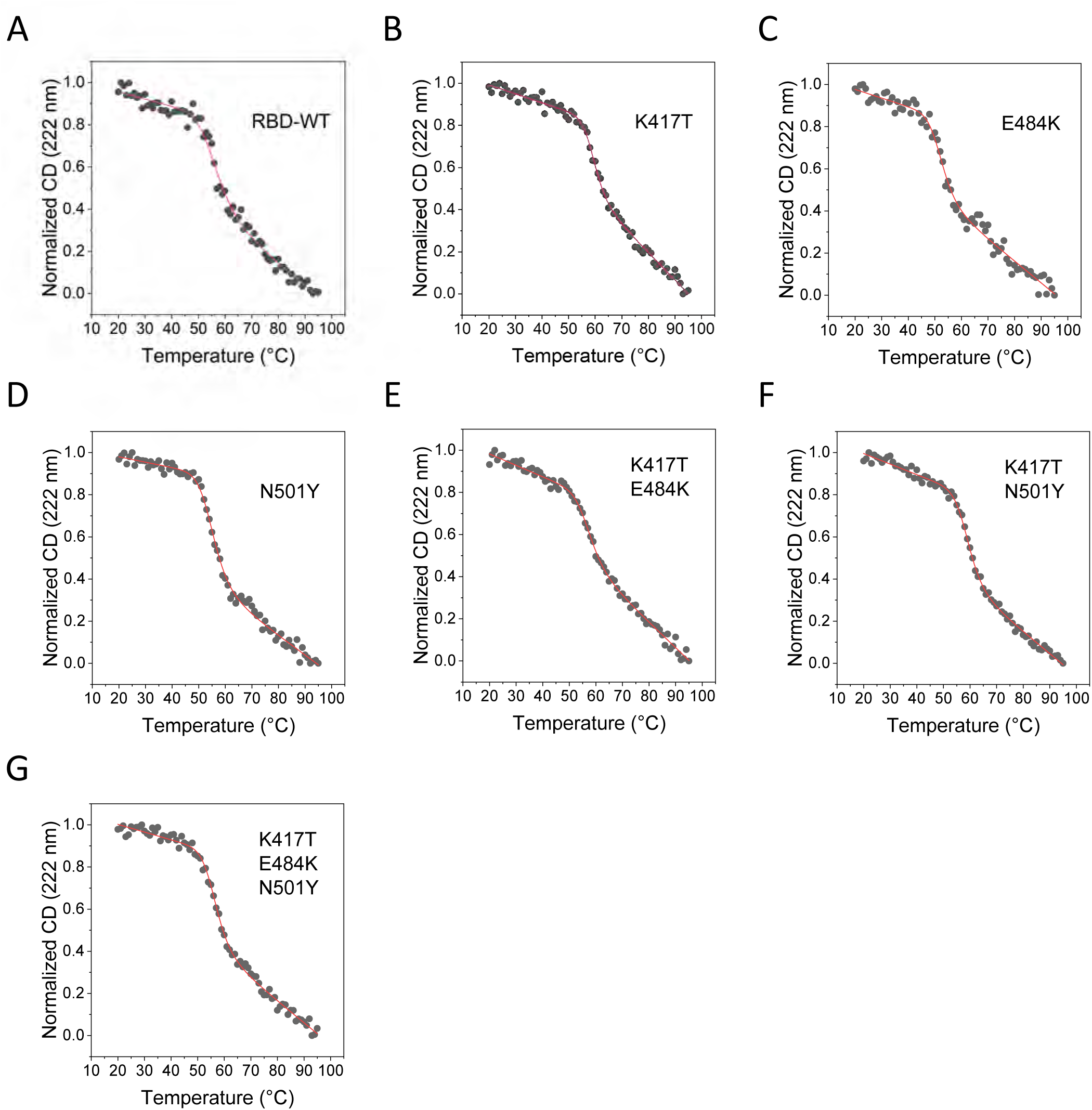
Comparison of thermal stability of WT RBD and its mutants measured using Far-UV CD spectroscopy. The data were fit to a two-state unfolding model. The individual data points are ellipticity values at 222 nm obtained at different temperatures and are represented as black circles. The red lines passing through the data points are the fit curves.

**Table 1.** T_m_ and ΔH values of RBD WT and its mutants determined using thermal denaturation melts.

| **RBD variant** | **$T_m$ (°C)** | **$\Delta T_m$ (°C)** | **$\Delta H_m$ (kcal/mol)** |
| --- | --- | --- | --- |
| RBD WT | $56.1 \pm 0.7$ | - | $-1.7 \pm 0.3$ |
| K417T | $59.3 \pm 0.2$ | $3.2 \pm 0.7$ | $-3.0 \pm 0.4$ |
| E484K | $52.3 \pm 0.5$ | $-3.8 \pm 0.9$ | $-2.0 \pm 0.3$ |
| N501Y | $55.1 \pm 0.3$ | $-1.0 \pm 0.8$ | $-2.1 \pm 0.1$ |
| K417T/E484K | $57.2 \pm 0.5$ | $1.1 \pm 0.9$ | $-2.2 \pm 0.3$ |
| K417T/N501Y | $59.1 \pm 0.2$ | $3.0 \pm 0.7$ | $-2.6 \pm 0.1$ |
| K417T/E484K/N501Y | $56.5 \pm 0.2$ | $0.4 \pm 0.7$ | $-2.4 \pm 0.2$ |

To obtain the true equilibrium thermodynamic stability of WT and mutant RBDs, chemical denaturation with urea was performed. The chemical denaturation melts were recorded using fluorescence spectroscopy. Chemical denaturation melts using CD were not used to calculate the stability of the proteins as the data was noisy below 220 nm in the presence of high concentrations of urea. β-sheet and random coil structures show maximum CD below 220 nm, and RBD is mainly composed of these secondary structures.^*30*^ Figures 4A-4G show representative urea denaturation melts, and Table 2 lists the fit parameters determined from three independent batches of protein expression. The chemical denaturation of RBD in urea using fluorescence was found to be completely reversible. The free energy change on unfolding (ΔG^0^_unf_) for the WT RBD was 8.1 ± 0.3 kcal/mol. Like the thermal denaturation melts, K417T mutation was found to stabilize and E484K mutation was found to destabilize the protein with ΔG^0^_unf_ values of 9.8 ± 0.3 and 7.0 ± 0.2 kcal/mol respectively. N501Y mutation was also found to destabilize with a ΔG^0^_unf_ value of 7.1 ± 0.2 kcal/mol. The double mutants that carried K417T mutation were more stable than the WT protein, with ΔG^0^_unf_ values of 8.7 ± 0.2 and 8.4 ± 0.2 kcal/mol for K417T/E484K and K417T/N501Y mutants, respectively. The triple mutant on the other hand had the least stability with ΔG^0^_unf_ value of 6.7 ± 0.2 kcal/mol, possibly showing the cooperative effect of two deleterious mutations E484K and N501Y on protein stability.

**Figure 4.**
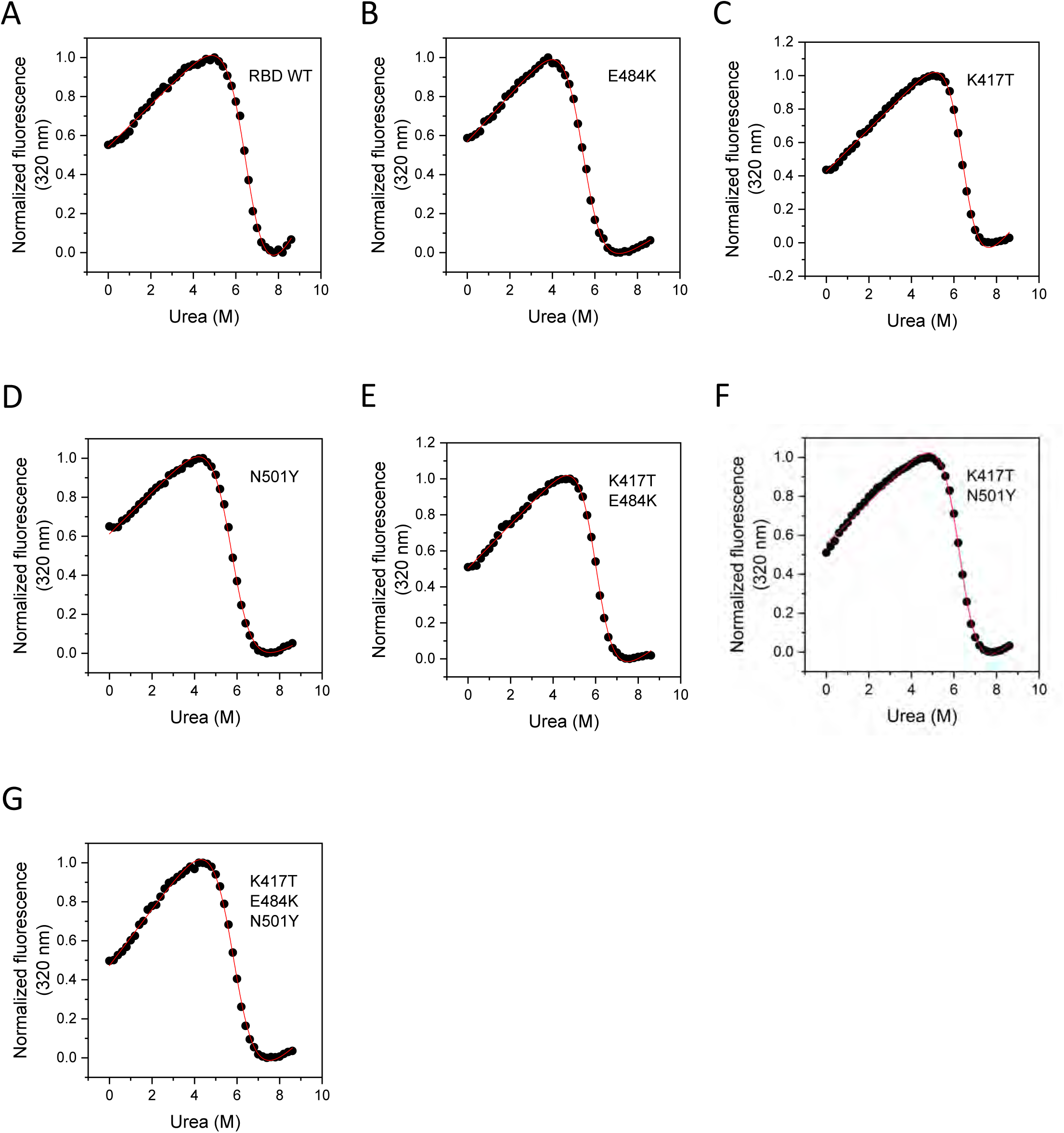
Comparison of protein stability of WT RBD and its mutants using chemical denaturation. Protein stability was measured using fluorescence spectroscopy following urea denaturation. Fluorescence intensity at 320 nm obtained at varying urea concentrations is represented as black circles. The fits passing through the data points are represented as red lines.

**Table 2.** ΔG^0^_unf_ and m-values determined from chemical denaturation melts of RBD WT and its mutants.

| **RBD variant** | **$\Delta G^0_{unf}$**<br>**(kcal/mol)** | **$\Delta\Delta G^0_{unf}$ (kcal/mol)** | **m-value**<br>**(kcal/mol/M [urea])** |
| --- | --- | --- | --- |
| RBD WT | $8.1 \pm 0.3$ | - | $-1.2 \pm 0.1$ |
| K417T | $9.8 \pm 0.3$ | $1.7 \pm 0.4$ | $-1.5 \pm 0.1$ |
| E484K | $7.0 \pm 0.2$ | $-1.1 \pm 0.4$ | $-1.3 \pm 0.1$ |
| N501Y | $7.1 \pm 0.2$ | $-1.0 \pm 0.4$ | $-1.2 \pm 0.1$ |
| K417T/E484K | $8.7 \pm 0.2$ | $0.6 \pm 0.4$ | $-1.5 \pm 0.1$ |
| K417T/N501Y | $8.4 \pm 0.2$ | $0.3 \pm 0.4$ | $-1.4 \pm 0.1$ |
| K417T/E484K/N501Y | $6.7 \pm 0.2$ | $-1.4 \pm 0.4$ | $-1.2 \pm 0.1$ |

### N501Y mutation enhances binding affinity towards ACE2

SARS-CoV-2 relies on the binding of RBD to ACE2 receptor on the host cells to enter the host and is thus an important parameter that determines virus infectivity and transmission. The mutations that increase the binding affinity towards ACE2 have a selective advantage and can naturally be selected during virus evolution. To determine which of the mutations provide the virus with selective advantage for improved ACE2 binding, binding affinity of the mutants towards ACE2 was determined using isothermal titration calorimetry (ITC) and compared it with the binding affinity of WT protein. Figures 5A-5G show representative ITC thermograms, and Table 3 lists the average fit parameters from three independent batches of protein expression. All proteins interacted with ACE2 in a stoichiometric ratio of 1:1. WT RBD interacts with ACE2 with a dissociation constant (K_d_) of 10.0 ± 3.1 nM. Of the three mutations, N501Y improved the binding affinity with a K_d_ value of 3.4 ± 0.4 nM (ΔK_d_ = 6.9 ± 3.1 nM). K417T and E484K mutations decreased the binding affinity with K_d_ values of 32.5 ± 8.9 and 51.5 ± 8.2 nM respectively. The presence of N501Y mutation in the double and the triple mutants (K417T/N501Y with K_d_ value of 8.8 ± 3.5 nM and K417T/E484K/N501Y with K_d_ value of 1.6 ± 1.5 nM) improved the binding affinity as compared to the variants that do not contain N501Y mutation (K417T with K_d_ value of 32.5 ± 8.9 nM and K417T/E484K with K_d_ value of 26.0 ± 3.3 nM). These results clearly suggest the importance of N501Y mutation in improving the binding affinity of the variants towards ACE2.

**Figure 5.**
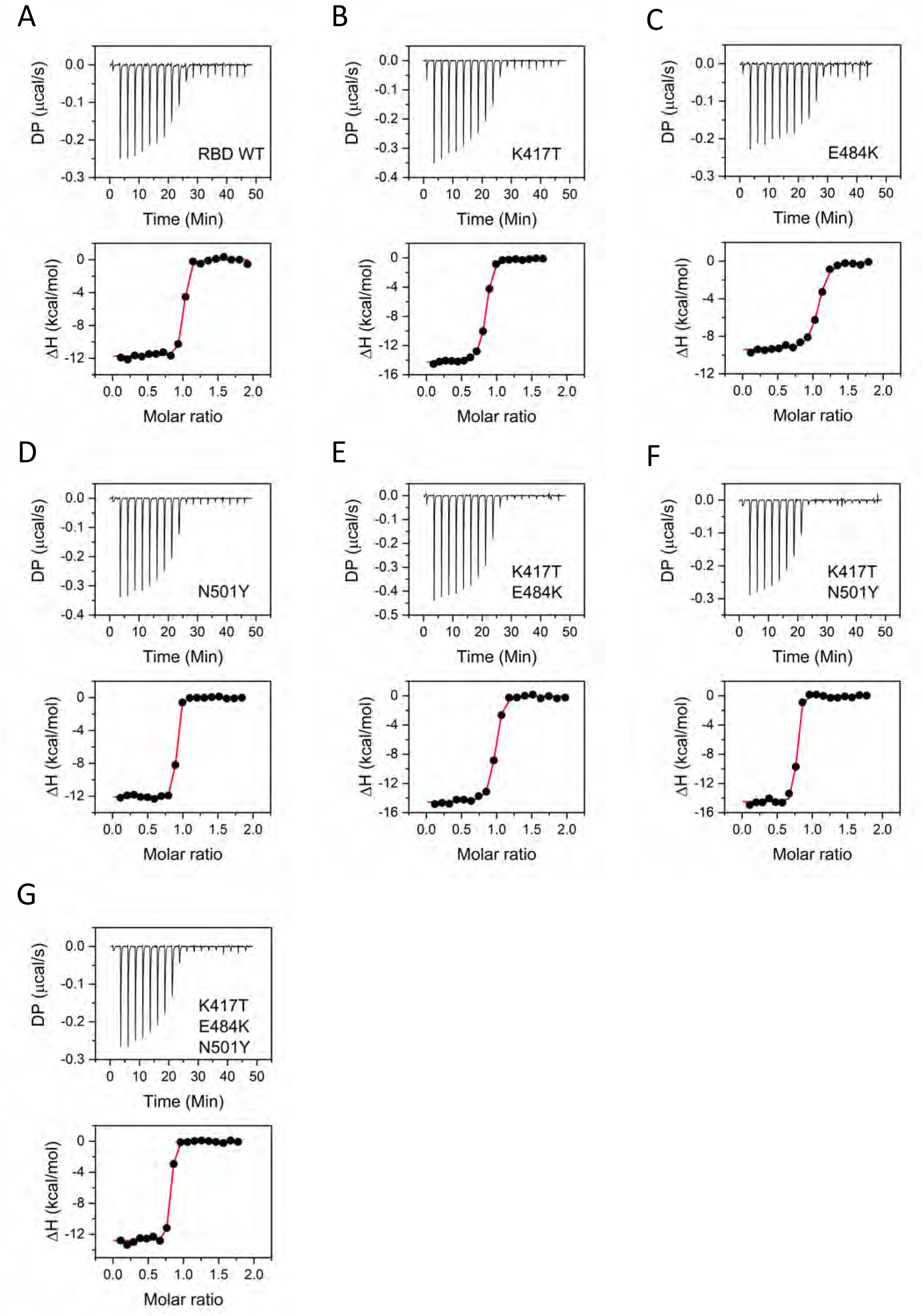
Binding of WT RBD and its mutants to ACE2 studied using ITC. The top panels show the raw thermograms. The bottom panels show the integrated heat at each injection in black circles and the fits passing through the data points are shown as red lines.

**Table 3.** Binding parameters for RBD WT and its mutants interaction with ACE2 determined from ITC.

| **RBD variant** | **K<sub>d</sub> (nM)** | **N** | **ΔH**<br>**(kcal/mol)** | **ΔG**<br>**(kcal/mol)** | **-TΔS**<br>**(kcal/mol)** |
| --- | --- | --- | --- | --- | --- |
| RBD WT | 10.0 ± 3.1 | 1.0 ± 0.1 | -11.8 ± 0.2 | -10.7 ± 0.2 | 1.0 ± 0.3 |
| K417T | 32.5 ± 8.9 | 0.9 ± 0.1 | -14.7 ± 0.4 | -10.1 ± 0.2 | 4.7 ± 0.4 |
| E484K | 51.5 ± 8.2 | 1.0 ± 0.1 | -9.5 ± 0.2 | -9.8 ± 0.1 | -0.3 ± 0.2 |
| N501Y | 3.4 ± 0.4 | 0.9 ± 0.1 | -11.8 ± 0.2 | -11.4 ± 0.1 | 0.5 ± 0.2 |
| K417T/E484K | 26.0 ± 3.3 | 0.9 ± 0.1 | -14.3 ± 0.4 | -10.2 ± 0.1 | 4.1 ± 0.4 |
| K417T/N501Y | 8.8 ± 3.5 | 0.8 ± 0.1 | -14.3 ± 0.3 | -10.9 ± 0.2 | 3.5 ± 0.3 |
| K417T/E484K/N501Y | 1.6 ± 1.5 | 0.9 ± 0.1 | -10.9 ± 0.2 | -11.8 ± 0.5 | -0.9 ± 0.6 |

### K417T mutation determines escape from Class 1 antibodies

The escape from neutralizing antibodies is one of the major factors that govern the natural selection of variants. The host antibody response towards the SARS-CoV-2 is varied with different antibodies produced against a range of viral epitopes, out of which only a few are neutralizing. RBD is the major target against which neutralizing antibodies are produced.^*22, 31–33*^ These neutralizing antibodies can be divided into different classes based on their site of interaction with RBD and consequently the mechanism of neutralization.^*22, 31*^ Majorly, two classes of antibodies (Class 1 and Class 2) bind to the site on RBD which is also the site of interaction with ACE2. Apart from that, Class 3 antibodies also share some region of the binding site with ACE2 (Figures 6A-6F). Although Class 4 antibodies are less in number, they do not block the ACE2 site, and the three mutations are far away in protein structure from the epitope region of Class 4 antibodies. Most of the RBD mutations in the emerging variants are present in the receptor binding motif (RBM), which is the region that interacts with ACE2. Thus, the three mutations are more likely to have an impact on the binding with Class 1 and Class 2 antibodies. Since Class 3 antibodies also partially occupy the ACE2 binding site, they can compete with the RBD mutants for binding to ACE2. To understand the emergence of mutations and the role of antibody escape in natural selection of mutants, we tested the binding of RBD mutants against Class 1, 2 and 3 antibodies.

**Figure 6.**
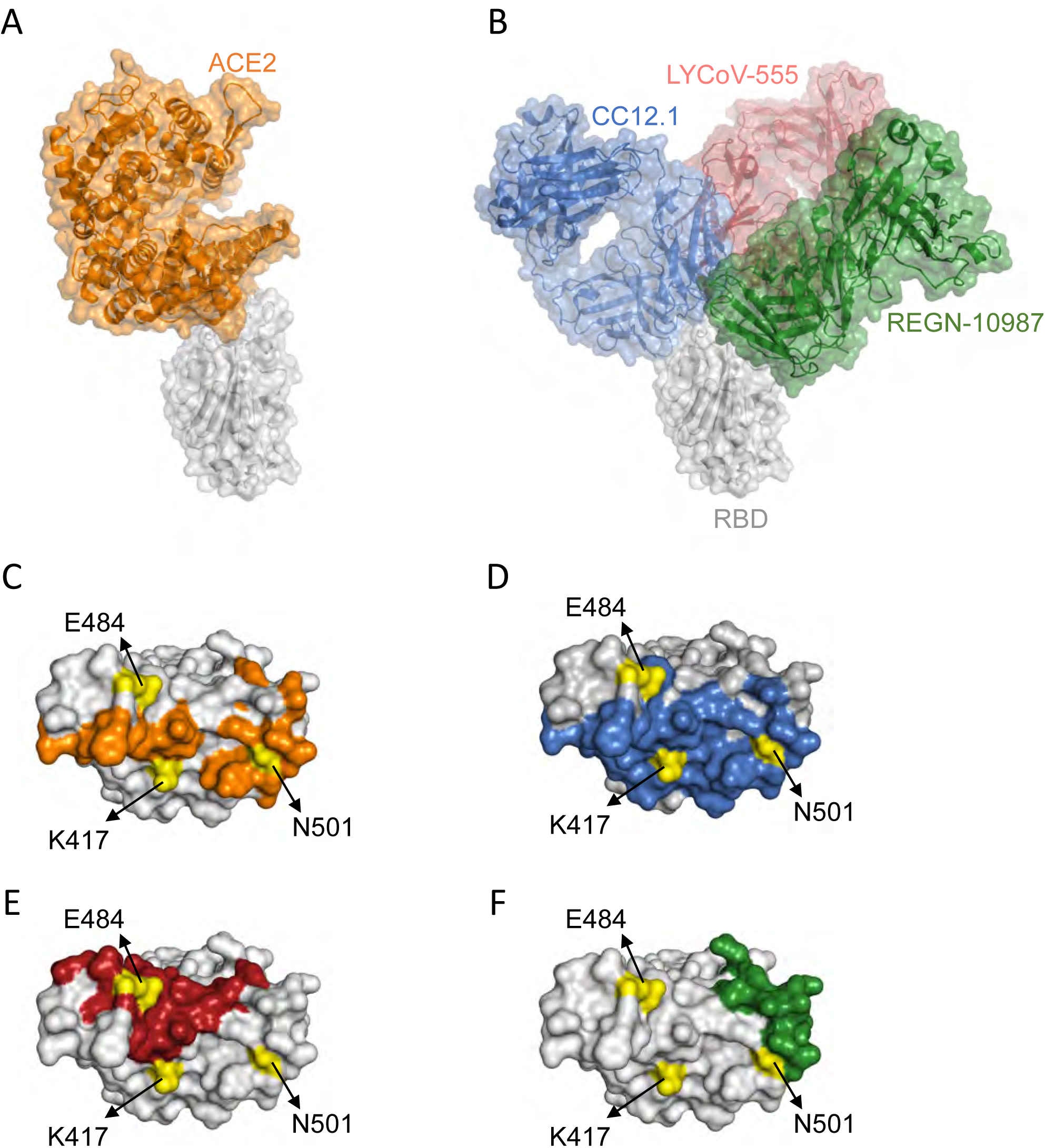
Interaction of SARS-CoV-2 RBD with ACE2 and neutralizing antibodies. (A) Protein structures showing interaction of RBD (gray) with ACE2 (orange) (Protein Data Bank (PDB) ID: 6m0j.pdb). (B) Protein structures showing interaction of RBD with a Class 1 antibody Fab, CC12.1 (blue) (PDB ID: 6xc2.pdb), a Class 2 antibody Fab, LYCoV-555 (red) (PDB ID: 7kmg.pdb), and a Class 3 antibody Fab, REGN-10987 (green) (PDB ID: 6xdg.pdb). (C-F) The RBD residues that interact with ACE2 (orange), CC12.1 (blue), LYCoV-555 (red) and REGN-10987 (green). The position of the three RBD amino acid residues are shown in yellow.

Most Class 1 antibodies represent a similar binding mode to RBD because they are encoded by the common VH3-53 gene segment and have short H3 CDR.^*34*^ We have used CC12.1 as a representative example of Class 1 antibodies and tested its interaction with WT and mutant RBDs. Figures 7A-7G show representative ITC thermograms, and Table 4 lists the average parameters obtained from three independent batches of protein expression. CC12.1 interacts with WT RBD with K_d_ value of 23.9 ± 5.7 nM. Out of the single site mutants, K417T decreased the binding affinity substantially with a K_d_ value of 301 ± 52 nM. The other single site mutants E484K and N501Y did not show escape from CC12.1 binding with K_d_ values of 1.7 ± 4.6 nM and 53.7 ± 12.1 nM respectively. All the other double and triple mutants that carried the K417T mutation also showed escape from CC12.1 binding with higher K_d_ values (429 ± 266 nM, 553 ± 273 nM and 402 ± 29 nM for K417T/E484K, K417T/N501Y and K417T/E484K/N501Y respectively). These results suggest that the escape from Class 1 antibodies is conferred predominantly by the K417T mutation.

**Figure 7.**
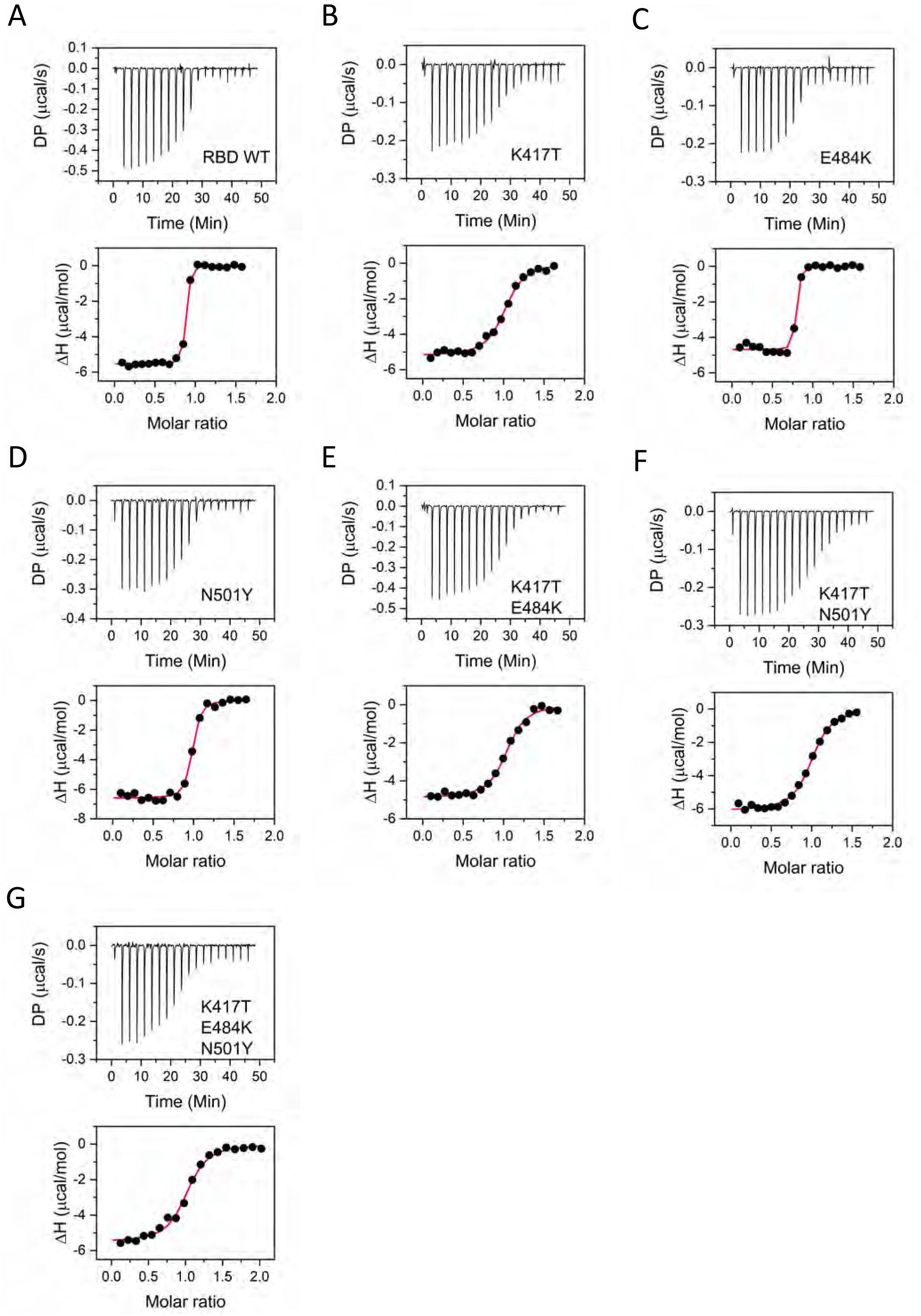
Binding of WT RBD and its mutants to CC12.1 ScFv studied using ITC. The top panels show the raw thermograms. The bottom panels show the integrated heat at each injection in black circles and the fits passing through the data points are shown as red lines.

**Table 4.** Binding parameters for RBD WT and its mutants interaction with CC12.1 ScFv determined from ITC.

| **RBD variant** | **K<sub>d</sub> (nM)** | **N** | **ΔH**<br>**(kcal/mol)** | **ΔG**<br>**(kcal/mol)** | **-TΔS**<br>**(kcal/mol)** |
| --- | --- | --- | --- | --- | --- |
| RBD WT | 23.9 ± 5.7 | 0.9 ± 0.1 | -5.2 ± 0.5 | -10.2 ± 0.1 | -5.1 ± 0.5 |
| K417T | 301 ± 52 | 1.0 ± 0.1 | -5.2 ± 0.1 | -8.8 ± 0.1 | -3.6 ± 0.1 |
| E484K | 1.7 ± 4.6 | 0.8 ± 0.1 | -5.5 ± 0.1 | -11.8 ± 1.6 | -6.2 ± 1.6 |
| N501Y | 53.7 ± 12.1 | 0.9 ± 0.1 | -6.4 ± 0.3 | -9.8 ± 0.1 | -3.4 ± 0.3 |
| K417T/E484K | 429 ± 266 | 1.0 ± 0.1 | -4.8 ± 0.1 | -8.6 ± 0.4 | -3.8 ± 0.4 |
| K417T/N501Y | 553 ± 273 | 1.0 ± 0.1 | -6.3 ± 0.3 | -8.4 ± 0.3 | -2.1 ± 0.4 |
| K417T/E484K/N501Y | 402 ± 29 | 1.0 ± 0.1 | -5.5 ± 0.1 | -8.6 ± 0.1 | -3.1 ± 0.1 |

### E484K mutation determines escape from Class 2 antibodies

Class 2 antibodies differ from Class 1 antibodies in the mode of interaction with RBD (Figure 6). Their epitope differs from that of Class 1 antibodies, and they can recognize RBD in both up and down conformation. One advantage of Class 2 antibodies is that they can recognize adjacent RBDs in the spike trimer and once bound they can lock RBDs in down conformation thereby restricting binding to ACE2.^*35*^ This difference in the mode of interaction with RBD is thought to be because of longer H3 CDR in Class 2 antibodies compared to Class 1 antibodies. LY-CoV555 antibody is a Class 2 antibody which was part of the Eli Lilly’s antibody cocktail authorized for emergency use in USA by FDA.^*36*^ We have used LY-CoV555 as a representative example of Class 2 antibodies and determined its binding to WT and mutant RBDs. Figures 8A-8G show representative thermograms, and Table 5 lists the average fit parameters obtained from three independent batches of protein expression. WT RBD interacts with LY-CoV555 with a K_d_ value of 3.8 ± 1.9 nM. N501Y mutant does not show any difference in the binding affinity with K_d_ value of 4.7 ± 0.4 nM. K417T mutant shows an increased K_d_ value of 32.8 ± 21.6 nM. Consistent with this data, the double mutant (K417T/N501Y) showed higher binding affinity with a K_d_ value of 1.6 ± 1.0 nM. E484K mutant on the other hand failed to bind to the LY-CoV555 antibody. Other double and triple mutants carrying E484K mutations also failed to bind to LY-CoV555, which clearly supports the role of E484K in the Class 2 antibody escape of the variants.

**Figure 8.**
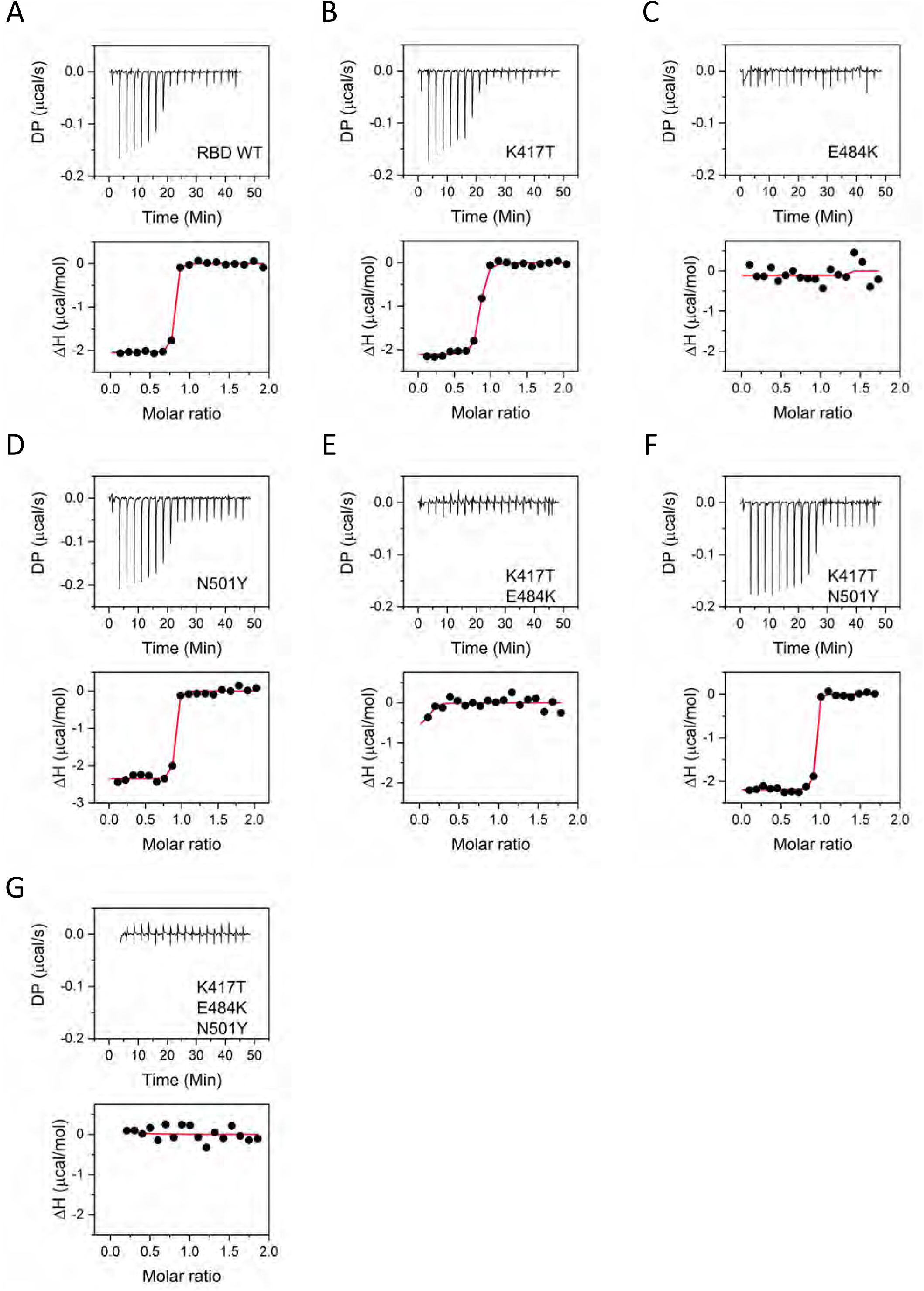
Binding of WT RBD and its mutants to LY-CoV555 ScFv studied using ITC. The top panels show the raw thermograms. The bottom panels show the integrated heat at each injection in black circles and the fits passing through the data points are shown as red lines.

**Table 5.**
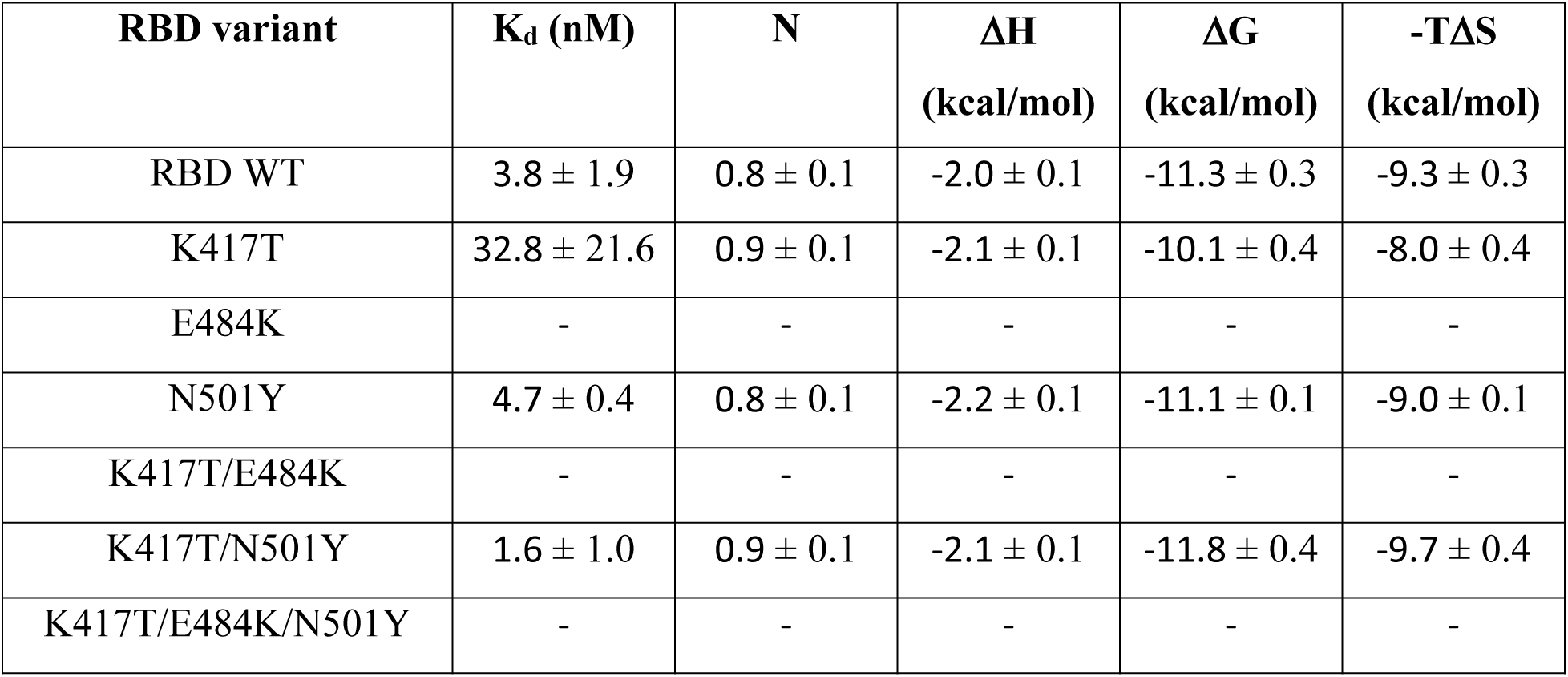
Binding parameters for RBD WT and its mutants interaction with LYCoV-555 ScFv determined from ITC.

### None of the mutants escape Class 3 antibodies

Class 3 antibodies do not block the ACE2 binding site and bind to the region of RBD that is adjacent to RBM (Figure 6). The three mutations K417T, E484K, and N501Y are thus not likely to affect the binding to Class 3 antibodies. We have used REGN-10987 as a representative example of Class 3 antibodies. REGN-10987 antibody was part of the Regeneron’s antibody cocktail authorized for emergency use in USA by FDA.^*36*^ Figures 9A-9G show representative ITC thermograms, and Table 6 lists the average fit parameters obtained from three independent batches of protein expression. WT protein interacts with REGN-10987 with a K_d_ value of 34.3 ± 8.1 nM. All the single site mutations K417T, E484K and N501Y did not show any major change in the binding affinity towards REGN-10987. Even the double and the triple mutants did not show any appreciable change and the K_d_ value for REGN-10987 interaction was between 5 -36 nM for all the variants. These results indicate that the three mutations do not escape Class 3 antibodies, and REGN-10987 will still be able to neutralize the variants containing these three mutations, unless the other mutations confer escape from Class 3 antibodies.

**Figure 9.**
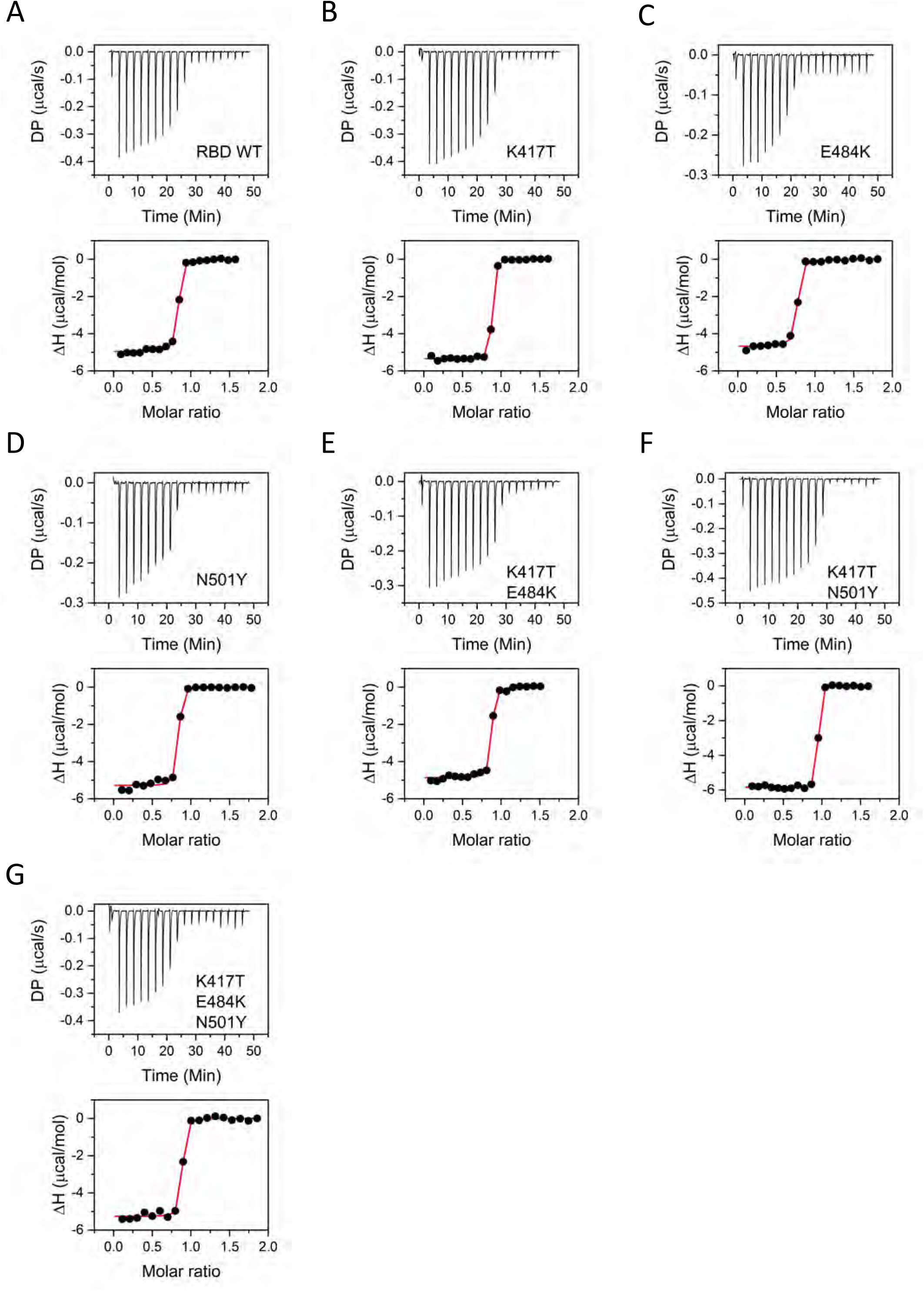
Binding of WT RBD and its mutants to REGN-10987 ScFv studied using ITC. The top panels show the raw thermograms. The bottom panels show the integrated heat at each injection in black circles and the fits passing through the data points are shown as red lines.

**Table 6.** Binding parameters for RBD WT and its mutants binding to REGN-10987 ScFv determined from ITC.

| **RBD variant** | **K<sub>d</sub> (nM)** | **N** | **ΔH**<br>**(kcal/mol)** | **ΔG**<br>**(kcal/mol)** | **-TΔS**<br>**(kcal/mol)** |
| --- | --- | --- | --- | --- | --- |
| RBD WT | 34.3 ± 8.1 | 0.8 ± 0.1 | -5.0 ± 0.1 | -10.0 ± 0.1 | -5.1 ± 0.2 |
| K417T | 10.6 ± 2.3 | 0.9 ± 0.1 | -5.3 ± 0.1 | -10.7 ± 0.1 | -5.4 ± 0.2 |
| E484K | 35.6 ± 11.1 | 0.7 ± 0.1 | -4.7 ± 0.1 | -10.0 ± 0.2 | -5.3 ± 0.2 |
| N501Y | 18.3 ± 7.4 | 0.8 ± 0.1 | -5.3 ± 0.1 | -10.4 ± 0.2 | -5.1 ± 0.3 |
| K417T/E484K | 15.8 ± 5.4 | 0.8 ± 0.1 | -4.9 ± 0.1 | -10.5 ± 0.2 | -5.6 ± 0.2 |
| K417T/N501Y | 4.6 ± 2.8 | 0.9 ± 0.1 | -5.8 ± 0.1 | -11.2 ± 0.4 | -5.3 ± 0.4 |
| K417T/E484K/N501Y | 14.1 ± 7.8 | 0.8 ± 0.1 | -5.3 ± 0.1 | -10.5 ± 0.3 | -5.3 ± 0.3 |

## DISCUSSION

The continued emergence of variants over a small timeframe means adaptive evolution of virus in the human host.^*37, 38*^ The adaptive evolution allows virus to counter the host immune response and at the same time maintains viral fitness. The three RBD mutations K417T, E484K and N501Y appear in multiple lineages, suggesting their convergent evolution where similar mutations appear in different lineages independently. The high frequency of occurrence of these mutations points to the positive selection of these mutations. Convergent evolution is thought to be a product of positive selection where advantageous mutations accumulate and become prevalent.^*39*^ The accumulation of distinct mutations that bring different selective advantages towards viral survival is a strategy that maximizes the virus fitness in its environment. Here we have delineated the role of individual mutations and their contribution to virus fitness in terms of stability, expression, ACE2 binding, and antibody escape. ACE2 binding is an important parameter that governs the natural selection of variants with high affinity interactions contributing to improved viral fitness.^*30, 40*^ Antibody selection pressure also impacts the emergence of variants with the escape variants better adapted to prevail.^*41–44*^ Apart from these parameters, protein stability also plays an important role in deciding the evolutionary path of a protein.^*45, 46*^ In the case of variants containing multiple mutations, one mutation can complement the existence of other mutation. The epistatic effects of mutations also need to be kept in mind as the virus evolves to accumulate multiple mutations. In the evolutionary history of the virus, certain mutations can only occur after a particular mutational event has occurred.^*47*^ This allows acceptance of even those mutations that have a deleterious effect individually. The results presented in this study suggest that each mutation serves a distinct purpose and coming together of these mutations improve the collective virus fitness over the effects of individual mutations.

Positively charged K417 residue in SARS-CoV-2 RBD has been shown to be beneficial towards ACE2 binding by forming electrostatic interactions with the negatively charged residues in the N-terminal helix of ACE2.^*48*^ K417 residue has been an intensive subject of mutations and K417N and K417T mutations have appeared with high frequencies in the emerging variants. K417T and K417N mutations rank 4^th^ and 6^th^ respectively on the list of mutations in RBD with highest frequency of occurrence (https://www.gisaid.org/hcov19-mutation-dashboard/). Change of residue from lysine to threonine has a penalty on the ACE2 binding affinity, which is not surprising as lysine residue provides a beneficial effect towards ACE2 binding. This means that the improved ACE2 binding is not a parameter which supports the positive selection of K417T mutation. The beneficial effect of K417T mutation is however seen in all the other aspects we examined. K417T mutation improved the expression of RBD and this effect was also observed for double mutants and the triple mutant carrying K417T mutation. K417T also increased the stability of RBD relative to the WT protein. It stabilized RBD by about 2 kcal/mol in chemical denaturation experiments (Table 2). Thermal denaturation melts also showed substantial increase in T_m_ by about 3°C (Table 1). Therefore, the role of K417T mutation is to provide stability to the variants which in turn helps in higher expression of the protein. Low expression levels of the double mutant E484K/N501Y reiterate the fact that K417T is needed for efficient protein expression carrying destabilizing mutations E484K and N501Y. Another interesting point in support of natural selection of K417T comes from the observation that although E484K and N501Y have been very frequent mutations that provide significant advantage to the virus, they do not appear together very frequently in evolving lineages unless accompanied by K417T/K417N mutation. Apart from increasing stability and expression, K417T mutation also contributes to escape from Class 1 antibodies. Substitutions at 417 position have been shown to be driven by antibody selection pressure, where incubation of the virus with Class 1 antibodies drive mutation at K417 position to E/N/T.^*49*^ In the CC12.1 antibody-RBD complex structure (Figure 10A), K417 residue lies in the middle of the interaction surface and interacts with D97 residue in CC12.1 heavy chain. Substitution to threonine breaks this ionic interaction and destabilize the RBD-antibody interface. Antibody escape potential of K417T can be an important contributing factor to its positive selection as Class 1 antibodies represent antibody class that are most commonly elicited in response to viral infection. These arguments provide a solid support for the positive selection of K417T mutation which improves virus fitness by increasing protein stability, enhanced protein expression and the ability to escape Class 1 antibodies even though ACE2 binding affinity is slightly compromised.

**Figure 10.**
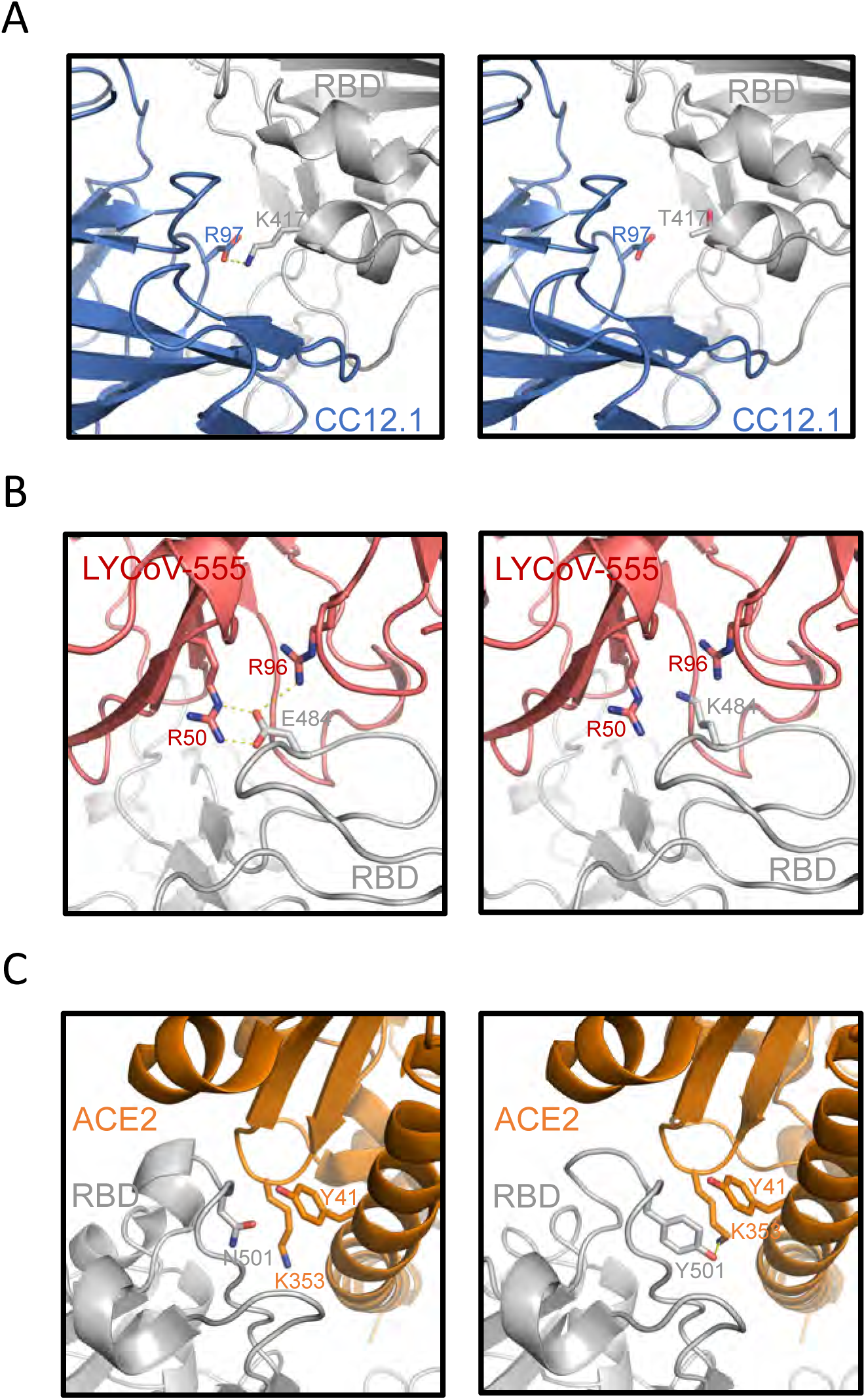
Comparison of wild type (left panel) and mutated (right panel) RBDs interaction with ACE2 and neutralizing antibodies. (A) Interaction of wild-type RBD (gray) lysine residue at 417 position (K417) with arginine residue at 97 (R97) position in the heavy chain variable region of CC12.1 antibody (blue) (PDB ID: 6xc2.pdb). Mutation of lysine to threonine at 417 position (K417T) in RBD disrupts this electrostatic interaction (K417T RBD structure from 7nx8.pdb was aligned with 6xc2.pdb). (B) Interaction of wild-type RBD (gray) glutamate residue at 484 position (E484) with arginine residues at 50 position (R50) in heavy chain and 96 position (R96) on the light chain variable region of LYCoV-555 antibody (PDB ID: 7kmg.pdb). Mutation of glutamate to lysine at 484 position (E484K) disrupts ionic interactions with variable domain arginine residues in LYCoV-555 antibody. Mutation was created using PyMOL mutagenesis wizard. (C) Wild-type RBD (gray) asparagine at 501 position (N501) shows weak interactions with lysine residue at 353 position (K353) and tyrosine residue at 41 position (Y41) of ACE2 (orange) (PDB ID: 6m0j.pdb). Mutation of asparagine to tyrosine at 501 position (N501Y) strengthens this interaction with K353 and Y41 residues of ACE2 (PDB ID: 7mjn.pdb).

E484K mutation has been one of the most commonly found mutations in SARS-CoV-2 evolution. This mutation ranks 3^rd^ in the GISAID list of most frequently occurring RBD mutations. E484K mutation can be found in a number of viral lineages like Beta, Gamma, Eta, Theta, Iota and Zeta.^*50*^ The presence of this mutation in independent viral lineages points towards convergent evolution where E484K mutation is independently selected in different variants. Independent emergence of E484K mutation in different viral populations incubated in the presence of antibodies stresses the importance of antibody selection pressure in natural selection of mutations.^*44*^ E484K mutation does not provide any selective advantage towards ACE2 binding as found previously by us and other groups and thus receptor binding is not a parameter that supports positive selection of E484K.^*30, 51, 52*^ E484K mutation also negatively impacts protein expression (Figure 1) and protein stability (Figures 3 and 4). As protein stability is an important factor that determines the inclusion of other mutations and also the persistence of mutations in viral lineages, a mutation that destabilizes the protein must have a strong reason for its so widespread and common selection in multiple lineages, which is provided by the antibody escape potential of E484K. Our results show that E484K escapes LYCoV-555, a Class 2 antibody. Amongst the three RBD mutations, only E484K lies on the interaction surface of LYCoV-555 (Figure 10B). The binding of WT RBD to LY-CoV555 antibody is substantially strengthened by the interaction of E484 with two arginine residues, R50 in the heavy chain and R96 in the light chain variable regions of LY-CoV555. E484K mutation is a non-conservative mutation and change to an oppositely charged lysine residue disrupts these electrostatic interactions and contributes to antibody escape. The importance of E484K mutation has been further highlighted by the observation that escape from Class 2 antibodies mainly contributes to escape from polyclonal plasma.^*53*^ These arguments present strong evidence for the positive selection of E484K mutation, and its convergent evolution conferred by its ability to escape neutralizing antibodies despite the penalty levied by decreased stability and expression. In the case of variants containing other mutations, the simultaneous presence of K417T mutation appears to mitigate the negative effects on stability and expression. The double mutant (K417T/E484K) and the triple mutant (K417T/E484K/N501Y) have expression levels comparable to that of the WT protein, and stability higher than the E484K mutant alone.

N501Y was amongst the first mutations that was identified and was the only RBD mutation in the Alpha variant, the first variant to be designated as the VOC.^*54, 55*^ Shortly after, Beta and Gamma variants emerged in different parts of the world, carrying N501Y mutation and were also designated as VOCs.^*56, 57*^ Omicron variants also contain N501Y mutation. N501Y is the 2^nd^ most frequently found mutation according to GISAID and points towards its positive selection. N501Y has been shown to improve the binding affinity towards ACE2 by us and other groups,^*30, 51, 58–61*^ and it is now established that the N501Y mutation increases the transmissibility and infectivity of the virus providing a selective advantage to virus fitness leading to its natural selection.^*62*^ E484K and K417T mutations individually do not provide any advantage towards ACE2 binding. The collective effect of three mutations still maintains higher binding affinity towards ACE2 compared to the WT protein. The effect of N501Y mutation on other parameters tested were not supportive of its positive selection. The expression and stability were slightly decreased and there was no significant escape towards any of the antibodies we tested although the presence of N501Y with K417T or E484K did not impact their antibody escape potential in either the double or the triple mutants.

## CONCLUSION

All the new variants that joined the list of VOCs after the Alpha variant contain multiple RBD mutations, and many mutations have been seen to evolve together in multiple variants. We examined the physical origins of such convergent evolution of mutations at three positions K417T, E484K, and N501Y in the primary structure of spike protein. The three RBD mutations perform very distinct and specific roles that contribute towards improving the virus fitness and builds the case for positive selection of these mutations (Table 7). K417T stabilizes the RBD and improves its expression. It also provides escape potential towards Class 1 antibodies. E484K provides escape potential towards Class 2 antibodies. N501Y improves ACE2 binding affinity important for virus infectivity and transmission. These single mutations, although bring advantages for virus fitness, they also bring deleterious effects, implying negative selection and decreasing the chances of these mutations occurring individually. K417T decreases the ACE2 binding affinity, whereas E484K and N501Y decreases stability and expression. However, the simultaneous presence of the three mutations mitigates the deleterious effects and enhances the chances of selection of mutations together. The collective effect of these mutations is far more advantageous for virus fitness than the individual mutations. Presence of multiple mutations thus improve the chances of survival of SARS-CoV-2 in the environment challenged by numerous selection pressures, mainly the antibody selection pressure. The three mutations we examined did not escape Class 3 antibodies. Any mutations that confer escape from Class 3 and Class 4 antibodies along with these three mutations could be worrisome.

**Table 7.** Comparison of protein expression, stability, binding to ACE2, and binding to neutralizing antibodies (Class 1-3) of WT RBD and its mutants.

| **RBD variant** | **Expression** | **T<sub>m</sub>**<br>**(°C)** | **ΔG**<br>**(kcal/mol)** | **K<sub>d</sub> (nM)**<br>**(ACE2**<br>**binding)** | **K<sub>d</sub> (nM)**<br>**(CC12.1**<br>**binding)**<br>**Class 1** | **K<sub>d</sub> (nM)**<br>**(LY-**<br>**CoV555**<br>**binding)**<br>**Class 2** | **K<sub>d</sub> (nM)**<br>**(REGN-**<br>**10987**<br>**binding)**<br>**Class 3** |
| --- | --- | --- | --- | --- | --- | --- | --- |
| RBD WT | 1.0 | 56.1 ± 0.7 | 8.1 ± 0.3 | 10.0 ± 3.1 | 23.9 ± 5.7 | 3.8 ± 1.9 | 34.3 ± 8.1 |
| K417T | 1.7 ± 0.3 | 59.3 ± 0.2 | 9.8 ± 0.3 | 32.5 ± 8.9 | 301 ± 52 | 32.8 ± 21.6 | 10.6 ± 2.3 |
| E484K | 0.4 ± 0.0 | 52.3 ± 0.5 | 7.0 ± 0.2 | 51.5 ± 8.2 | 1.7 ± 4.6 | No binding | 35.6 ± 11.1 |
| N501Y | 0.8 ± 0.0 | 55.1 ± 0.3 | 7.1 ± 0.2 | 3.4 ± 0.4 | 53.7 ± 12.1 | 4.7 ± 0.4 | 18.3 ± 7.4 |
| K417T/E484K | 1.1 ± 0.1 | 57.2 ± 0.5 | 8.7 ± 0.2 | 26.0 ± 3.3 | 429 ± 266 | No binding | 15.8 ± 5.4 |
| K417T/N501Y | 1.4 ± 0.2 | 59.1 ± 0.2 | 8.4 ± 0.2 | 8.8 ± 3.5 | 553 ± 273 | 1.6 ± 1.0 | 4.6 ± 2.8 |
| E484K/N501Y | 0.2 ± 0.0 | - |  | - | - | - | - |
| K417T/E484K<br>/N501Y | 1.1 ± 0.2 | 56.5 ± 0.2 | 6.7 ± 0.2 | 1.6 ± 1.5 | 402 ± 29 | No binding | 14.1 ± 7.8 |

## MATERIALS AND METHODS

### Cloning and expression of WT RBD and its mutants, ACE2 and antibodies in ScFv format

The genes encoding WT RBD, ACE2 and the antibodies (CC12.1, LY-CoV555 and REGN10987) in ScFv format were commercially obtained from Twist Biosciences carrying BamH1 and Xho1 restriction sites on the 5’ and 3’ end respectively. The genes ware introduced into the modified pcDNA3.4 TOPO mammalian expression vector using BamH1 and Xho1 site. The modified vector contained a human immunoglobulin heavy chain signal sequence, His-tag and SUMOstar protein. RBD mutants were created with site directed mutagenesis using mutagenic primers. Vectors containing the gene of interest were transfected transiently into Expi293 cells using polyethyleneimine (PEI) as the transfection agent. The cells were tested for protein expression 3 days post transfection. The protein expression levels were compared using SDS-PAGE followed by quantitative densitometric analysis of Coomassie stained protein bands using Image-J software. After 5 days of post transfection, the proteins were recovered from the culture supernatant for purification.

### Protein purification

The culture supernatant was filtered through 0.22 μm PVDF membrane filter and purified using Nickel-nitrilotriacetic acid (Ni-NTA) chromatography. The eluted protein was cleaved with SUMOstar protease to remove the SUMOstar and His-tag. The cleaved protein was again passed through the Nickel column to obtain the untagged protein in the flow-through. The proteins were exchanged into buffer containing 50 mM Sodium Phosphate, 20 mM NaCl, pH 7.0. All the experiments were performed in this buffer unless mentioned otherwise.

### Circular dichroism (CD) spectroscopy

Far-UV CD spectra of the proteins were collected at a protein concentration of 5 μM in buffer containing 10 mM Sodium Phosphate, 4 mM NaCl, pH 7.0, on an Applied Photophysics Chirascan Plus spectrometer. The spectra were collected in a wavelength range of 190 – 260 nm at an interval of 1 nm and averaging time of 2 s at each wavelength. Thermal denaturation melts were obtained at protein concentration of 20 μM. The temperature ramp rate was kept at 1°C/min. CD signal at 222 nm obtained at an interval of 1°C with averaging time of 2 s was plotted against temperature to obtain the denaturation curve. The data were fit to a two-state unfolding model using the equation^*29*^

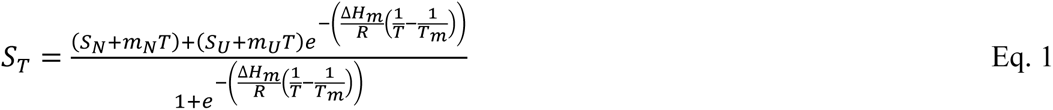

Where S_T_ is the measured ellipticity signal with respect to temperature T, S_N_ and S_U_ are the ellipticity signals corresponding to the native and unfolded baselines, m_N_ and m_U_ are the slopes of linear dependence of S_N_ and S_U_, ΔH_m_ is the enthalpy change at T_m_, R is the universal gas constant, and T is the absolute temperature in Kelvin. Errors on ΔT_m_ values were calculated using error propagation formulae.^*63*^

### Fluorescence spectroscopy

Fluorescence emission spectra of proteins were collected at protein concentration of 2 μM on a PTI Quantamaster fluorimeter. The samples were excited at 280 nm and emission spectra were collected in wavelength range of 300-400 nm. Data were acquired at an interval of 1 nm and averaging time of 1 s at each wavelength. Chemical denaturation melts using fluorescence spectroscopy were obtained with an automated titrator connected to an Applied Photophysics Chirascan Plus CD spectrometer equipped with a CCD detector for fluorescence measurements. Different urea concentrations ranging from 0 - 8.6 M at an interval of 0.2 M were achieved by mixing two end state solutions of native and denatured protein in a 1 cm path length cuvette. Urea stock concentration was determined by refractive index measurements.^*64*^ The equilibration time for mixing was kept as 5 min. The samples were excited at 280 nm and the change in fluorescence emission intensity at 320 nm with urea concentration was plotted to obtain the chemical denaturation curves. The raw data were fit to a two-state unfolding model using the equation^*65*^

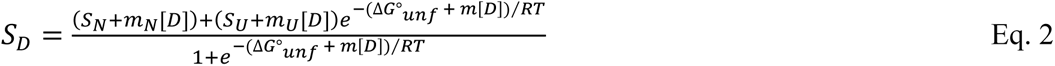

where S_D_ is the measured fluorescence intensity signal at 320 nm as a function of denaturant concentration D, S_N_ and S_U_ are the fluorescence signals corresponding to the native and unfolded baselines, m_N_ and m_U_ are the slopes of linear dependence of S_N_ and S_U_, ΔG°_unf_ is the free energy change for unfolding, R is the universal gas constant, and T is the absolute temperature in Kelvin. Errors on ΔΔG^0^_unf_ values were calculated using error propagation formulae.^*63*^

### Isothermal titration calorimetry

The interaction of WT RBD and the mutants with ACE2 and different antibodies were performed on MicroCal-PEAQ-ITC instrument from Malvern. For ACE2 - RBD interaction, 15 μM ACE2 was taken in the cell and 150 μM RBD or its mutants were taken in the syringe for titrations. For RBD-ScFv interactions, 25 - 40 µM RBD or its mutants were taken in the cell and 250 - 400 μM of ScFv of different antibodies were taken in the syringe. The titrations were performed as sets of 18 injections of 2 μl each with equilibration time of 150 s between injections. All ITC data were analyzed using MicroCal PEAQ-ITC Analysis Software from Malvern. Errors on ΔG and -TΔS were calculated using error propagation formulae.^*63*^

